# Genomic landscape of the DHA1 family in the newly emerged pathogen *Candida auris* and mapping substrate repertoire of the prominent member *Cau*Mdr1

**DOI:** 10.1101/2022.03.09.483563

**Authors:** Rosy Khatoon, Suman Sharma, Poonam Vishwakarma, Amandeep Saini, Parth Aggarwal, Andrew M. Lynn, Amresh Prakash, Rajendra Prasad, Atanu Banerjee

## Abstract

The last decade has witnessed the rise of extremely threatening healthcare-associated multidrug resistant *non-albicans Candida* (NAC) species, *Candida auris*. Thus, understanding the molecular basis of antifungal resistance has emerged as the single most important goal amongst the research community. Since besides target alterations, efflux mechanisms contribute maximally to antifungal resistance, it is imperative to investigate their contributions in this pathogen. Of note, within the Major Facilitator Superfamily (MFS) of efflux pumps, Drug/H+ antiporter family 1 (DHA1) has been established as a predominant contributor towards xenobiotic efflux. Our study provides a complete landscape of DHA1 transporters encoded in the genome of *C. auris*. This study identifies 14 DHA1 transporters encoded in the genome of the pathogen. We also construct deletion and heterologous overexpression strains for the most important DHA1 drug transporter, viz., *Cau*Mdr1 to map the spectrum of its substrates. While the knockout strain did not show any significant changes in the resistance patterns against the majority of the tested substrates, the ortholog when overexpressed in a minimal background *S. cerevisiae* strain, AD1-8u^-^, showed significant enhancement in the Minimum inhibitory concentrations (MICs) against a large panel of antifungal molecules. Altogether, the present study provides a comprehensive template for investigating the role of DHA1 members of *C. auris* in antifungal resistance mechanisms.

## INTRODUCTION

The recent emergence of multidrug resistant *C. auris* in clinical settings within a short period across five continents and 42 countries is a global health concern (Chakrabarti and Sood, 2021; Du et al., 2020). *C. auris* clusters into 4 major clades, with a recent report suggesting a possible fifth clade from Iran (Chow et al., 2019; Muñoz et al., 2018). Its multidrug resistant profile alongside the paucity of antifungal agents has prompted the search for alternative therapeutics (Frías-De-León et al., 2020). However, the success of the latter is dependent on a comprehensive understanding of the resistance mechanisms in *C. auris*. Recent assessments of resistance patterns suggest about 90 % of *C. auris* strains are resistant to fluconazole, 30 % towards amphotericin B, and approximately 5 % to echinocandins. Furthermore, about 41 % of the strains also display multidrug resistance (MDR) (Chakrabarti and Sood, 2021). Amongst the different clades of *C. auris*, clade I strains show maximal resistance, wherein 97 % of the isolates show resistance to fluconazole, 54 % show resistance to amphotericin B (AmB), and 49 % manifesting MDR (Chow et al., 2020). In terms of resistance towards AmB in clinical isolates, no direct mechanism was known until recently, when Rybak et al. revealed a mutation in *ERG6* as a contributing factor (Rybak et al., 2021). In the case of echinocandins, substitutions in the typical FKS1 hotspot region, viz., S639P, S639F, and S639Y are frequently observed (Chakrabarti and Sood, 2021; Chowdhary et al., 2018). Compared to echinocandins and polyenes, azole resistance mechanisms are much more elaborately described in *C. auris*. Drug target alterations, such as mutations in the *ERG11* gene resulting in amino acid substitutions such as Y132F, K143R, and F126L are frequently seen in azole-resistant *C. auris* isolates (Chakrabarti and Sood, 2021; Chow et al., 2020; Lockhart et al., 2016). Fluconazole exposure has also been shown to induce lanosterol 14-α-demethylase overexpression via an increase in *ERG11* copy numbers (Chowdhary et al., 2018). Recently Rybak and colleagues exploited experimental evolution experiments to show that serial fluconazole exposure can result in resistance in an otherwise susceptible isolate via accumulation of mutations in *TAC1B* (homolog of the well-known *C. albicans* transcriptional regulator *CaTAC1* (Coste et al., 2004)) and a corresponding increase in the expression of ATP binding cassette (ABC) drug transporter encoding gene *CDR1* (Rybak et al., 2020). Similar mutations were also identified in numerous clinical isolates and correction of one such mutation in a resistant isolate significantly compromised the fluconazole resistance (Rybak et al., 2020). Notwithstanding that certain studies also point towards the existence of Tac1-dependent and Cdr1-independent azole resistance mechanisms (Li et al., 2021), efflux mediated azole resistance seems a prominent case in *C. auris*. Notably, several azole-resistant clinical isolates show an upregulation of ABC transporter encoding genes either naturally or on drug exposure (Jenull et al., 2021; Rybak et al., 2019; Wasi et al., 2019; Zamith-Miranda et al., 2019). Similar observations were also made by Carolus et. al. with a susceptible isolate on *in vitro* exposure to azoles, polyenes, and echinocandins (Carolus et al., 2021). Of note, Rybak et. al also confirmed the role of the prominent member Cdr1 in azole resistance mechanism, by constructing deletion mutants which led compromised resistance in *C. auris* isolates to a major extent (Rybak et al., 2019). While a clinical correlation of ABC transporter encoding gene and azole resistance has been experimentally deduced, the same is yet unknown for the other major class of drug transporters in *Candida* species, the Major Facilitator superfamily (MFS) (Prasad et al., 2019, 2017). Within the MFS of fungi, the drug/H^+^ antiporter 1 family (DHA1) proteins have shown capability to recognize and export a wide variety of xenobiotic compounds, including drugs like azoles (Banerjee et al., 2021; Prasad et al., 2017). The major clinically relevant DHA1 transporter in *Candida* species is Mdr1 and gene encoding for its homolog also exists in *C. auris*, which shows increased transcript levels in resistant isolates (Bravo Ruiz and Lorenz, 2021; Jenull et al., 2021; Rybak et al., 2019). However, one study showed that deletion of the gene did not result in a major alteration in the MICs. Nonetheless, it is worth mentioning that there is a significant overlap between the substrate repertoire of DHA1 transporters and ABC transporters, and function of one protein might not be apparent in the presence of a large number of other such proteins (Banerjee et al., 2021). Further research is still in infancy and more studies with a larger panel of isolates are required to understand contributions of these important class of proteins to antifungal resistance. This becomes much more important with recent observations like overexpression of other DHA1 family members in azole-resistant *C. auris* isolates (Jenull et al., 2021; Zamith-Miranda et al., 2019). Given such evidence, the current study is aimed at identification of all the DHA1 transporter encoding gene as well as to map the substrates of the prominent transporter *Cau*Mdr1.

## MATERIALS AND METHODS

### DHA1 transporter identification

Representative sequences of DHA1 transporters were downloaded from TCDB and multiple sequence alignment was performed using in-house scripts as described before (Vishwakarma et al., 2018). This alignment is used to build an HMM profile using the hmmbuild program of HMMER 3 package with default settings (Eddy, 2009). The proteome file (generated by translating all the ORFs) of the *C. auris* reference strain B8441 present in the Candida genome database (CGD) (“C_auris_B8441_current_orf_trans_all.fasta.gz”) was used as the target database. To identify DHA1 sequences in this database, the generated HMM profile was used as a query to match against the proteome (target database) with the help of the “hmmsearch” function within the HMMER 3 package using the default settings.

### Identification of Protein domains

Protein sequences of all the predicted DHA1 transporters were retrieved from CGD and a combined FASTA file was submitted as an input to the DomainViz webserver (Schläpfer et al., 2021). Default parameters were utilized for the job.

### Multiple sequence alignment and clustering sequence identity matrix construction

Multiple sequence alignment (MSA) of the DHA1 protein sequences was constructed using PSI/TM-Coffee (http://tcoffee.crg.cat/apps/tcoffee/do:tmcoffee) (Floden et al., 2016). Within the homology search options, sequence type was set to “transmembrane” and UniRef100 was selected for homology extension. The downloaded MSA was also used to generate the percent identity matrix using the BioEdit sequence alignment editor. The resultant sequence identities were used to generate the clustering sequence identity matrix using the Seaborn data visualization library (https://seaborn.pydata.org/generated/seaborn.clustermap.html) in Python.

### Phylogenetic analysis of DHA1 transporters

Protein sequences of previously identified DHA1 transporters in *S. cerevisiae* and *C. albicans* (Dias and Sá-Correia, 2014; Sá-Correia et al., 2009) were retrieved from Saccharomyces Genome Database (SGD) and CGD, respectively. Next protein sequences from all three species were arranged in a FASTA file and used for the construction of MSA using PSI/TM-Coffee as discussed before. The MSA generated above was then utilized to generate the phylogenetic tree in MEGA-X (Kumar et al., 2018) using the Maximum likelihood method. The bootstrap consensus tree inferred from 1000 replicates was taken to represent the evolutionary history of the taxa analyzed.

### Materials

All routine chemicals used were purchased from SRL Pvt. Ltd., Mumbai, India. Fluconazole (FLC) Nile red (NR), clotrimazole (CTZ), itraconazole (ITR), ketoconazole (KTZ), miconazole (MCZ), voriconazole (VOR), anisomycin (ANI), cerulenin (CER), 4-Nitroquinoline 1-oxide (4-NQO), cycloheximide (CHX) and dimethyl sulfoxide (DMSO) were procured from Sigma-Aldrich Co. (St. Louis, MO). Nourseothricin Sulfate was purchased from Gold biotechnology Inc. (USA). Oligonucleotides were purchased from Bioserve Biotechnologies India Pvt. Limited.

### Strains and culture conditions

All the Yeast strains used in the study are listed in **Supplementary table 1**. The *C. auris* strain CBS10913T was obtained from the CBS-KNAW fungal culture collection of the Westerdijk Fungal Biodiversity Institute, Utrecht, Netherlands, and reported in our previous study (Wasi et al., 2019). The yeast strains were grown in YEPD medium (yeast extract, peptone, and dextrose) from HiMedia Laboratories, Mumbai India. The *S. cerevisiae* transformants were selected on a Synthetic Defined medium without uracil. The medium comprised of 0.67 % YNB (Yeast nitrogen base) without amino acids (Difco, Becton, Dickinson and Company, MD, USA), 0.2% dropout mix without uracil (Sigma-Aldrich Co.), and 2% glucose (HiMedia). Plasmids were maintained in the *Escherichia coli* Dh5α strain, cultured in Luria-Bertani medium (HiMedia Laboratories, Mumbai, India) with 100 μg/ml ampicillin (Amresco, Solon, USA).

### Construction of *MDR1* deletion mutant in CBS10913T

For the construction of deletion mutant, we used the fusion PCR-based deletion strategy. Herein, the ORF is replaced with NAT1 cassette, which codes for nourseothricin acetyltransferase and imparts resistance to nourseothricin. Briefly, 5’-and 3’-UTR region (nearly 650 bps) of the gene was amplified from genomic DNA of *C. auris* CBS10913T. The selection marker *NAT1* gene with FRT sites was also PCR amplified in two halves from plasmid pRK625. Both 5’-and 3’-UTR amplified products were fused to one-half each of the NAT1 gene fragments to generate a fusion product. The fused PCR products were then co-transformed into the CBS10913T strain using electroporation strategy and positive transformants were selected on YEPD agar medium containing 200 μg/ml nourseothricin. The deletion was confirmed by genomic DNA PCR followed by DNA sequencing. The primers used in *MDR1* deletion and confirmation are listed in **Supplementary table 2**.

### Molecular cloning of *MDR1* in *pABC3-GFP* and *pABC3-His* and overexpression in AD1-8u^-^

*MDR1* gene was amplified from the genomic DNA of *C. auris* CBS10913T strain using forward and reverse primers containing *Pac*I and *Not*I restriction sites respectively (**Supplementary table 2**). The amplicons were purified using the Monarch PCR clean-up kit (New England Biolabs, MA, USA). The purified fragments and the pABC3-*His* or *pABC3-GFP* vector were digested with *Pac*I and *Not*I restriction enzymes (New England Biolabs, MA, USA). Resultant fragments were then gel-purified using the Monarch DNA Gel extraction kit (New England Biolabs, MA, USA) and ligated with the T4 DNA ligase (New England Biolabs, MA, USA). Positive clones (*pABC3-MDR1-His* or pABC3-*MDR1-GFP*) were confirmed by restriction digestion, followed by DNA sequencing. The *pABC3-MDR1-His* or *pABC3-MDR1-GFP* plasmid was then digested with *Asc*I to liberate the transformation cassette, which was then gel-purified using the Monarch DNA Gel extraction kit (New England Biolabs, MA, USA). *S. cerevisiae* AD1-8u^-^ (Decottignies et al., 1998; Lamping et al., 2007) cells were then transformed with the purified cassette using the LiAc method. The transformants were selected on SD-Ura^-^ media. Positive colonies were identified using gene-specific PCR.

### Confocal microscopy

For microscopy, exponential phase cells of AD-*Cau*Mdr1-GFP strain were washed with 1X PBS buffer before being examined under Nikon confocal microscope (Model:-A1R HD 25) with a 60X oil immersion objective lens.

### Broth microdilution assay (BMD)

MIC_80_ for the tested antifungal substrates were determined according to the CLSI broth microdilution method using YEPD medium instead of RPMI (Tanabe et al., 2019). Briefly, a two-fold serial dilution of each compound was prepared in YEPD medium and incubated with logarithmic phase cells (~10^4^ cells/ml) at 30°C for two days. The MIC_80_ value for a particular compound was considered as the lowest concentration that inhibited growth by 80%.

### Nile red accumulation assay

Nile red accumulation was studied as described before (Redhu et al., 2018) using the BD Facs Lyric set up (Becton–Dickinson Immunocytometry systems, San Jose, CA, USA), and data analysis was performed using the BD FACSuite v1.3. Median fluorescence intensity (MFI) was determined and represented using histograms made in GraphPad Prism 9.

### Statistical analysis

The plots in the study were made using either MS-Excel or GraphPad Prism 9 (San Diego, CA) or the Seaborn package in Python. Data for the Nile red accumulation assay is represented as mean ± SD. Statistical analysis for this experiment was performed using the Unpaired T-test with Welch’s correction. Differences were considered statistically significant when *p*< 0.05 (*, **, *** and **** indicate *p* values **≤** 0.05, 0.01, 0.001, and 0.0001, respectively).

## RESULTS AND DISCUSSION

### Inventory of DHA1 transporters encoded in the genome of *C. auris*

To identify the DHA1 transporters encoded in the genome of *C. auris*, we exploited a method based on the Hidden Markov Model. Herein, we first downloaded all the sequences listed under the TCDB # 2.A.1.2, representing The Drug: H^+^ Antiporter-1 (12 Spanner) (DHA1) family and constructed an MSA using in-house scripts as reported previously (Vishwakarma et al., 2018). The MSA was then used to build an HMM profile which was then used as a query against the *C. auris* proteome file as elaborated in the methods section. E-value and score (we call it domain score) values for all the sequences above the default threshold in the output HMM file **(Supplementary file 1)** were then used to create a plot **(Figure 1A)**. Positive hits above the default threshold are then further filtered based on a cutoff defined from this plot. This cutoff is interpreted in terms of a sudden large drop in the domain score and a consequent increase in the E-value as reported in our previous publications (Vishwakarma et al., 2021; Wasi et al., 2019). As represented in **Figure 1A**, there is a significant drop in the domain score following sequence number 14^th^ implying that the top 14 sequences are true DHA1 transporters. To further confirm their status as MFS proteins, we exploited the DomainViz web server that uses PFAM and Prosite databases for domain searching and displays the extent of domain conservation along two dimensions: positionality and frequency of occurrence in the input protein sequences (Schläpfer et al., 2021). As evident from **Figure 1B**, MFS_1 (PF07690) is the most represented PFAM domain, with all the submitted sequences containing it, confirming all to be part of the MFS superfamily. Of note, some proteins show similarity also to the PFAM domain, Sugar_tr (PF00083) which includes sugar (and other) transporters and one representative B9J08_001213 also shows similarity to TRI12 (PF06609) denoting Fungal trichothecene efflux pump.

**Figure 1:**
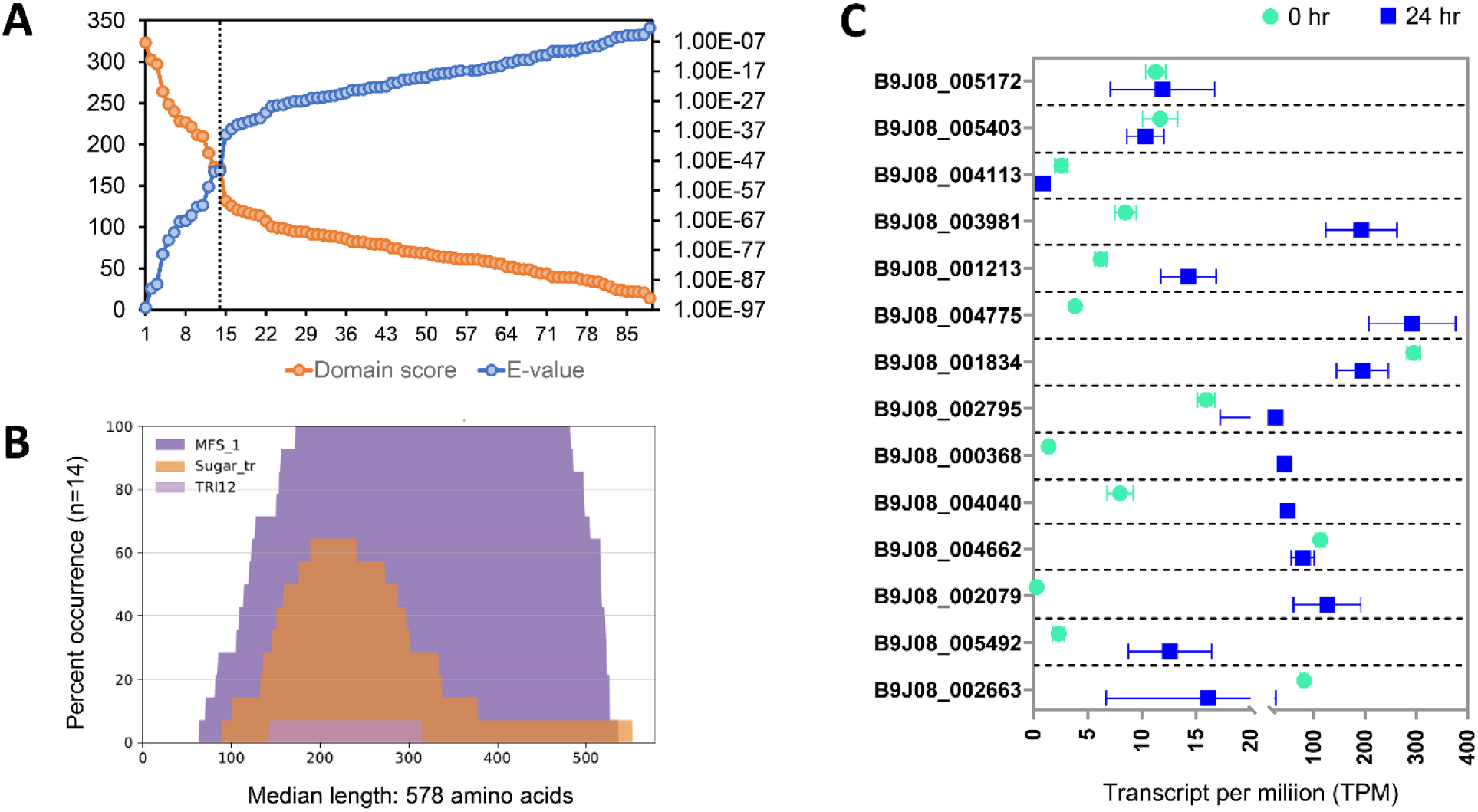
Identification of DHA1 transporters in *Candida auris*. **(A)** Plot of E-value and domain score of the sequences above the inclusion threshold value obtained after running the “hmmsearch” function with the *C. auris* proteome. **(B)** Distribution of PFAM domains in the identified DHA1 protein sequences as predicted using DomainViz. **(C)** Expression status of all the identified DHA1 transporter encoding genes is represented as Transcript per million (TPM). The data is obtained from the FungiDB database which reports the expression data from Kean et al. (Kean et al., 2018). The data depicts gene expression changes in the clinical isolate NCPF8973 between 0 hr planktonic and 24hr biofilm stages.

### Identification of orthologs of the identified DHA1 transporters in *S. cerevisiae* and *C. albicans*

To have a greater understanding of the identified protein’s homology and their plausible functions, we identified each candidate’s orthologs in *S. cerevisiae* and *C. albicans*,respectively from CGD. The results are presented in **Table 1**. Interestingly, as per the findings, proteins encoded by B9J08_003981 and B9J08_004113 are predicted orthologs of the prominent DHA1 multidrug transporter Mdr1 of *C. albicans* (*Ca*Mdr1). Furthermore, Rybak et al. have reported another protein encoded by B9J08_000368 to share high homology with *Ca*Mdr1 (Rybak et al., 2019). Notably, while B9J08_003981 and B9J08_000368 products both have *S. cerevisiae* Flr1 as their ortholog, *Sc*Yhk8 is the predicted ortholog for the protein encoded by B9J08_004113. Of note, Flr1 is an established multi-xenobiotic efflux pump showing the overlap in substrate spectrum with *Ca*Mdr1 (Alarco et al., 1997; Sá-Correia et al., 2009). On the other hand, literature regarding Yhk8 remains minimal, with certain studies pointing to its differentially expressed status in association with reduced azole susceptibility (Barker et al., 2003). Given the situation, we constructed an MSA with the protein sequences of all three probable candidates and *Ca*Mdr1. It was observed that while the B9J08_003981 protein sequence shows ~60% identity, both B9J08_000368 and B9J08_004113 only have approximately ~22% identity with *Ca*Mdr1, emphasizing B9J08_003981 as the true ortholog. Qdr1-3 are another set of DHA1 transporters that are implicated in resistance towards multiple xenobiotics/drugs like quinidine, barban, etc., in *S. cerevisiae* (Nunes et al., 2001; Tenreiro et al., 2005; Vargas et al., 2004). In the case of *Candida* species, only the *Candida glabrata Cg*Qdr2 has been reported to confer imidazole drug resistance (Costa et al., 2013), while the *C. albicans* counterpart is only involved in biofilm development and virulence (Shah et al., 2014). Interestingly, 4 plausible Qdr transporter encoding genes are present in *C. auris*: B9J08_005492, B9J08_002663, B9J08_005172, and B9J08_005403. Interestingly, both B9J08_005172 and B9J08_005403 are predicted to be orthologs of Qdr3, and B9J08_005492 as the ortholog of Qdr1 in *S. cerevisiae* and *C. albicans*. Notably, B9J08_002663 is predicted to be the ortholog of Qdr2 and Aqr1 in *C. albicans* and *S. cerevisiae*, respectively. *Sc*Aqr1 is known for conferring resistance towards compounds like quinidine and amino acid excretion (Tenreiro et al., 2002; Velasco et al., 2004). In our dataset, we also noticed an ortholog of *Ca*Flu1 (B9J08_002795), which is another gene that has been implicated in azole resistance, albeit marginally (Calabrese et al., 2000). Secondly, *Ca*Flu1 has been also identified as an exporter of Salivary antifungal peptide, Histatin 5 (Li et al., 2013). Interestingly, the predicted ortholog of B9J08_002795 in *S. cerevisiae* is Tpo1, which is a polyamine transporter showing high homology to *Ca*Flu1 (Li et al., 2013). Orthologs of the polyamine transporter *Ca*Tpo3 and *Ca*Tpo4 are also present in *C. auris*., viz. B9J08_004775 and B9J08_001834, respectively, with the former also showing homology to *Sc*Tpo2. Interestingly, certain Tpo members in *C. glabrata* have been reported to confer azole resistance too (Pais et al., 2016). The *C. auris* genome also encodes for a putative N-acetylglucosamine (GlcNAc) transporter (B9J08_001213), an ortholog of *Ca*NAG4 which is also implicated in cycloheximide resistance (Yamada-Okabe and Yamada-Okabe, 2002). Furthermore, two putative Hol transporters (orthologs of *Sc*Hol1), B9J08_004040 and B9J08_004662 are also encoded in *C. auris* genome. The *S. cerevisiae* transporters are involved in cation transport (Gaber et al., 1990). Finally, one ortholog of the *S. cerevisiae* dityrosine transporter Dtr1 (Felder et al., 2002) was also identified in our dataset, encoded by B9J08_002079.

**Table 1:** DHA1 transporter encoding genes identified in the genome of *Candida auris* and their orthologs in *Saccharomyces cerevisiae* and *Candida albicans*.

| **Sequence ID** | **Ortholog in *C. albicans*** | **Ortholog in *S. cerevisiae*** |
| --- | --- | --- |
| B9J08_001213 | C6_04620C_A/NAG4 | TPO3 |
| B9J08_004775 | C1_08790W_A/TPO3 | TPO2 |
| B9J08_002795 | C7_01520W_A/FLU1 | TPO1 |
| B9J08_003981 | C6_03170C_A/MDR1 | FLR1 |
| B9J08_000368 | C3_03440C_A | FLR1 |
| B9J08_001834 | CR_03920C_A/TPO4 | TPO4 |
| B9J08_005172 | C6_01290C_A/QDR3 | QDR3 |
| B9J08_005403 | C6_01290C_A/QDR3 | QDR3 |
| B9J08_005492 | CR_04210C_A/QDR1 | QDR1 |
| B9J08_002663 | C3_05570W_A/QDR2 | AQR1 |
| B9J08_002079 | CR_04620C_A | DTR1 |
| B9J08_004113 | C6_03170C_A/MDR1 | YHK8 |
| B9J08_004040 | C1_10200C_A | HOL1 |
| B9J08_004662 | C1_01780C_A/HOL4 | HOL1 |

To have a fair understanding of their expression status, we mined the transcriptomic dataset reported in FungiDB (Basenko et al., 2018). The RNA-Seq based transcriptome data present in the database originates from the study by Kean et al. which focused on temporally developing *C. auris* biofilms (Kean et al., 2018). We studied the expression changes of the predicted DHA1 transporter encoding genes in 0 hr planktonic and 24 hr biofilm stage of a clinical isolate, NCPF8973, originally isolated from a wound swab. As evident from the Transcript per million (TPM) data provided in **Figure 1C**, several predicted DHA1 transporter encoding genes show significant upregulation in the biofilm stage. Since *C. auris* biofilms are known to manifest significant antifungal resistance (Sherry et al., 2017), increased transcript levels in the biofilm stage imply a plausible role of these genes in the antifungal resistance mechanism. The significantly upregulated candidates include: B9J08_003981 (ortholog of *Ca*Mdr1), B9J08_001213 (ortholog of *Ca*Nag4/*Sc*Tpo3), B9J08_004775 (ortholog of *Ca*Tpo3/*Sc*Tpo2), B9J08_000368 (ortholog of *Sc*Flr1), B9J08_004040 (ortholog of *Sc*Hol1), B9J08_002079 (ortholog of *Sc*Dtr1) and B9J08_005492 (ortholog of *Ca*Qdr1/*Sc*Qdr1). Of note, orthologs of the majority of these proteins in *S. cerevisiae* and/or *Candida* species are previously implicated in conferring resistance towards antifungal drugs/compounds (Redhu et al., 2016; Sá-Correia et al., 2009).

### Sequence-based clustering and phylogenetic studies with identified DHA1 transporters of *C. auris*

Since sequence identity is often responsible for shared substrate preferences, we next sought to determine sequence relatedness between the identified DHA1 transporters. To achieve the same, we constructed a clustering sequence identity matrix using MSA of the protein sequences. As shown in **Figure 2A**, one major and another minor cluster is evident from the matrix. The minor cluster represents the 2 putative Qdr3 orthologs, i.e., B9J08_005172 and B9J08_005403 as reported in the previous section. Their separate clustering is due to the high sequence identity. The major cluster further comprises of two groups containing 7 and 5 members each, the former group showing moderately more sequence identity amongst themselves compared to the other group. The 5-member cluster includes one subcluster including B9J08_004040 (ortholog of *Sc*Hol1) and B9J08_004662 (ortholog of *Sc*Hol1 and *Ca*Hol4), and another subcluster with B9J08_005492 (ortholog of *Sc*Qdr1 and *Ca*Qdr1), B9J08_002663 (ortholog of *Ca*Qdr2 and *Sc*Aqr1) as part of one group, and B9J08_002079 (ortholog of *Sc*Dtr1) as an outgroup. Interestingly, there is a fair extent of substrate overlap between *Sc*Aqr1, *Sc*Qdr1, and *Sc*Dtr1 (Sá-Correia et al., 2009). The 7-member cluster contains B9J08_004113 (ortholog of *Sc*Yhk8) as an outgroup and the rest of the proteins, which are orthologs of *Sc*Flr1/*Ca*Mdr1 or the Tpo group of polyamine transporter of *S. cerevisiae* are included as a subcluster.

**Figure 2:**
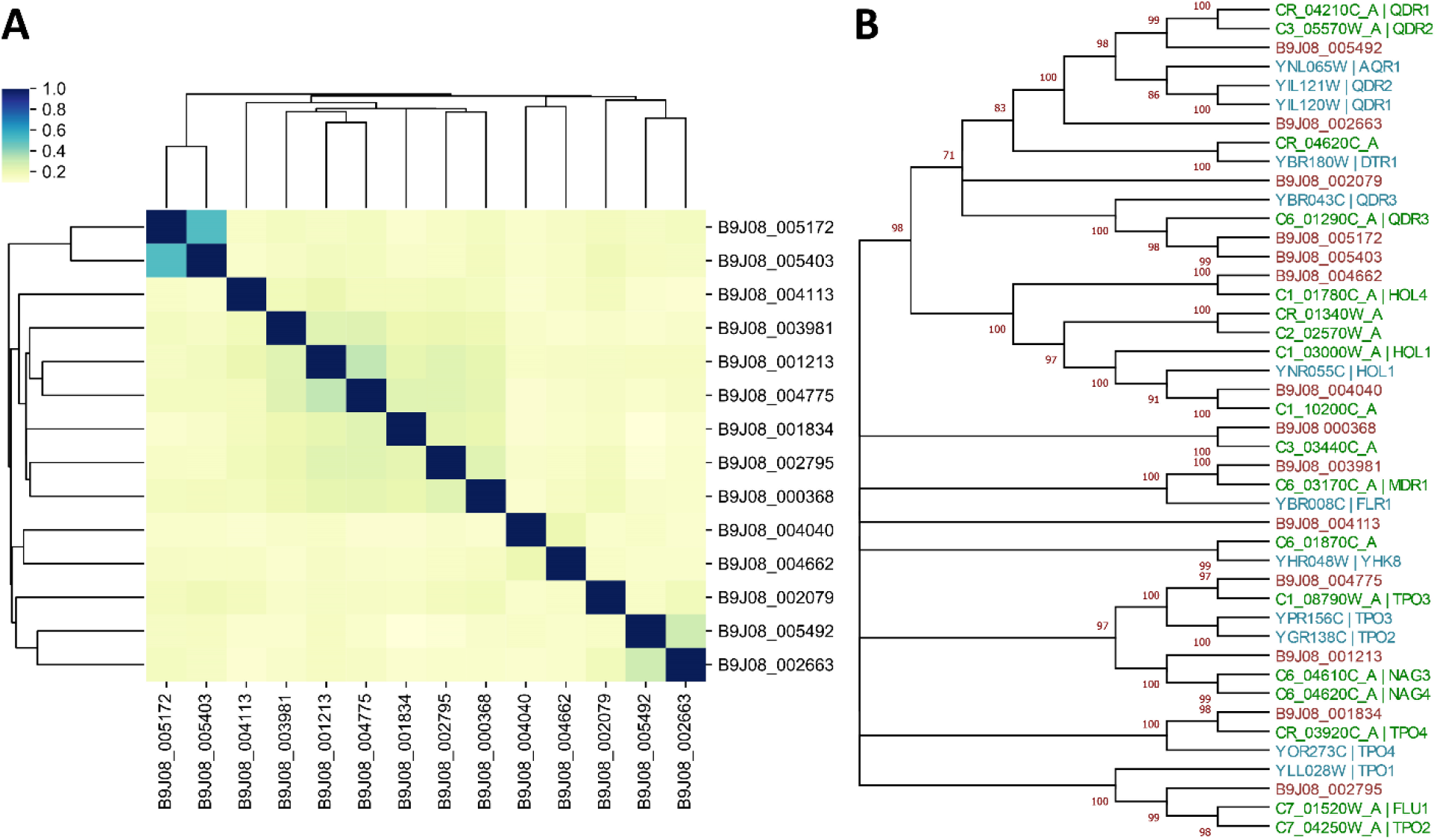
Sequence relatedness and phylogenetic analysis of DHA1 transporters. **(A)** Clustering sequence identity matrix of the predicted DHA1 protein sequences prepared using Seaborn library in Python. The input identity values for the plot were calculated from MSA generated using the BioEdit tool. **(B)** Phylogenetic tree of the DHA1 transporters from *S. cerevisiae* (Sá-Correia et al., 2009), *C. albicans* (Dias and Sá-Correia, 2014) and *C. auris* (this study) constructed using the Maximum Likelihood method and Jones et al. w/freq. model. The bootstrap consensus tree inferred from 1000 replicates is taken to represent the evolutionary history of the taxa analyzed. The percentage of replicate trees in which the associated taxa clustered together in the bootstrap test (1000 replicates) are shown next to the branches. Initial tree(s) for the heuristic search were obtained automatically by applying Neighbor-Join and BioNJ algorithms to a matrix of pairwise distances estimated using the JTT model, and then selecting the topology with a superior log likelihood value.

To have a further insight into the evolutionary relationship between the DHA1 transporters of *S. cerevisiae*, *C. albicans*, and *C. auris*, we constructed a phylogenetic tree using the maximum likelihood method. As presented in **Figure 2B**, it could be seen that the DHA1 family is distributed into several clades, the largest comprising of two clusters, namely the Aqr1/Dtr1/Qdr and the Hol. Notably, the *Ca*Mdr1/*Sc*Flr1 can be seen as a distinct clade along with B9J08_003981, further supporting its role as the true ortholog of *Ca*Mdr1. Interestingly, the Tpo group of polyamine transporters is part of several distinct clades. In accordance with our orthology assessment, *Ca*Flu1, *Ca*Tpo2, *Sc*Tpo1, and B9J08_002795 are part of one clade, and CaTpo4, ScTpo4, and B9J08_001834 part of another. Of note, another major clade is evident which comprises 2 clusters: ScTpo2/Tpo3, CaTpo3, and B9J08_004775 as one and other containing Nag transporters of *C. albicans* along with B9J08_001213. This observation implies that B9J08_001213 encodes by all chances, an N-acetylglucosamine (GlcNAc) transporter despite being a predicted ortholog of ScTpo3. Together, the identity-based clustering and the phylogenetic analysis strengthen our overall prediction of the orthologs and will thus, help prediction of their functional roles and substrate specificities.

### Construction of deletion mutant of *C. auris* ortholog of *Ca*Mdr1

It is evident from the previous analysis that B9J08_003981 is the true ortholog of *MDR1* gene. To understand its impact on the resistance profile of *C. auris* towards antifungal compounds, we deleted the gene from *C. auris* strain CBS10913T using the fusion PCR-based methodology as elaborated in the methods section. The deletion mutant was then exposed to different antifungal drugs and molecules, the majority of which are previously reported to be substrates of *Ca*Mdr1 **(Table 2)** (Redhu et al., 2016). While Rybak et al. also constructed deletion mutant of *Ca*Mdr1 in a clinical isolate their panel of tested substrates was restricted to the azole drugs only. In line with previous observations (Rybak et al., 2019), we did not notice any significant reduction in the MI_80_ values against most of the tested molecules except a modest reduction in the case of Itraconazole **(Table 2)**. This can be mainly attributed to the fact that most of the tested drugs/xenobiotics are established substrates of other efflux pumps in *S. cerevisiae* and *Candida* species (Prasad et al., 2015; Redhu et al., 2016; Sá-Correia et al., 2009).

**Table 2:** MIC_80_ values (μg/ml) determined for various antifungal drugs/xenobiotics in liquid culture as described in the methods section. The concentration (μg/ml) at which growth was inhibited by 80% remained the same for all the tested xenobiotics in three independent experiments.

| Antifungals/<br>Xenobiotics | MIC<sub>80</sub> (µg/ml) |  |  |  |
| --- | --- | --- | --- | --- |
|  | CBS10913T | *MDR1Δ* | AD1-8u<sup>-</sup> | AD-*CauMdr1* |
| **Fluconazole** | 16-32 | 16-32 | 0.5 | 64-128 |
| **Ketoconazole** | 0.06 | 0.06 | 0.002 | 0.004 |
| **Miconazole** | 0.25 | 0.25 | <0.001 | <0.001 |
| **Voriconazole** | 0.25 | 0.25 | <0.03 | 1 |
| **Clotrimazole** | 0.25 | 0.25 | <0.002 | 0.008 |
| **Itraconazole** | 0.016 | 0.004 | <0.001 | <0.001 |
| **Cerulenin** | 1 | 1 | 0.25 | 16 |
| **4-NQO** | 0.125 | 0.125 | 0.06 | 2 |
| **Cycloheximide** | > 8 | > 8 | <0.016 | 2 |
| **Anisomycin** | 16 | 16 | <0.25 | 8 |

### Overexpression of *Cau*Mdr1 in a heterologous *S. cerevisiae* strain to map the substrate repertoire

Since we were unable to identify the substrates recognized by *Cau*Mdr1 using the deletion mutant, we overexpressed the protein in an established *S. cerevisiae* host system, AD1-8u^-^ which is devoid of seven prominent antifungal drug transporters (Lamping et al., 2007). In addition, the strain also carries the *pdr1-3* gain of function mutation which hyperinduces the *PDR5* promoter under whose regulation we place the *CaMDR1* gene. This system has been extensively utilized to map the substrates of fungal efflux pumps, including *Ca*Cdr1 and *Ca*Mdr1 (Banerjee et al., 2020; Lamping et al., 2007; Redhu et al., 2018). We overexpressed *Cau*Mdr1 as both His and GFP-tagged fusion proteins. The construction of the AD-*Cau*Mdr1-GFP strain helped us inspect the expression and membrane localization of the *Cau*Mdr1 transporter through confocal microscopy. The confocal images shown in **Figure 3** demonstrate the localization of the transporter to the cell membrane as expected. However, to study the resistance attributes, we used the AD-*Cau*Mdr1-His strain to exclude any influence of the large GFP tag in the phenotypes. MIC_80_ values provided in **Table 2**, show significantly enhanced resistance levels of AD-*Cau*Mdr1-His strain compared to the host strain against several tested substrates including fluconazole, voriconazole, clotrimazole, cerulenin, cycloheximide, 4-NQO, and anisomycin. Notably, in the case of ketoconazole, the increase is only 2-fold for the AD-*Cau*Mdr1-His strain relative to the host AD1-8u^-^. It is important to mention that we could not determine the range of differential susceptibility (if at all) in the case of miconazole and itraconazole because both host and AD-*Cau*Mdr1-His strains were susceptible to these drugs even up to a very low concentration (0.001 μg/ml). Interestingly, when the resistance profile of AD-*Cau*Mdr1-His strain is compared with that of AD-*Ca*Mdr1-His strain as reported in our previous studies (Redhu et al., 2018), there is an evident differential pattern of resistance. For instance, AD-*Cau*Mdr1-His strain is more resistant towards fluconazole than the AD-*Ca*Mdr1-His strain. It will be of interest to explore whether the differences in the type of residues forming the drug binding pocket (only 60% sequence identity) contributes to such differential resistance patterns, or it is due to different levels of expression/membrane localization. Nonetheless, it is quite clear that *Cau*Mdr1 is capable of imparting resistance towards a wide spectrum of antifungal drugs and xenobiotics, most likely through its efflux function as witnessed in other *Candida* species too (Prasad et al., 2017).

**Figure 3:**
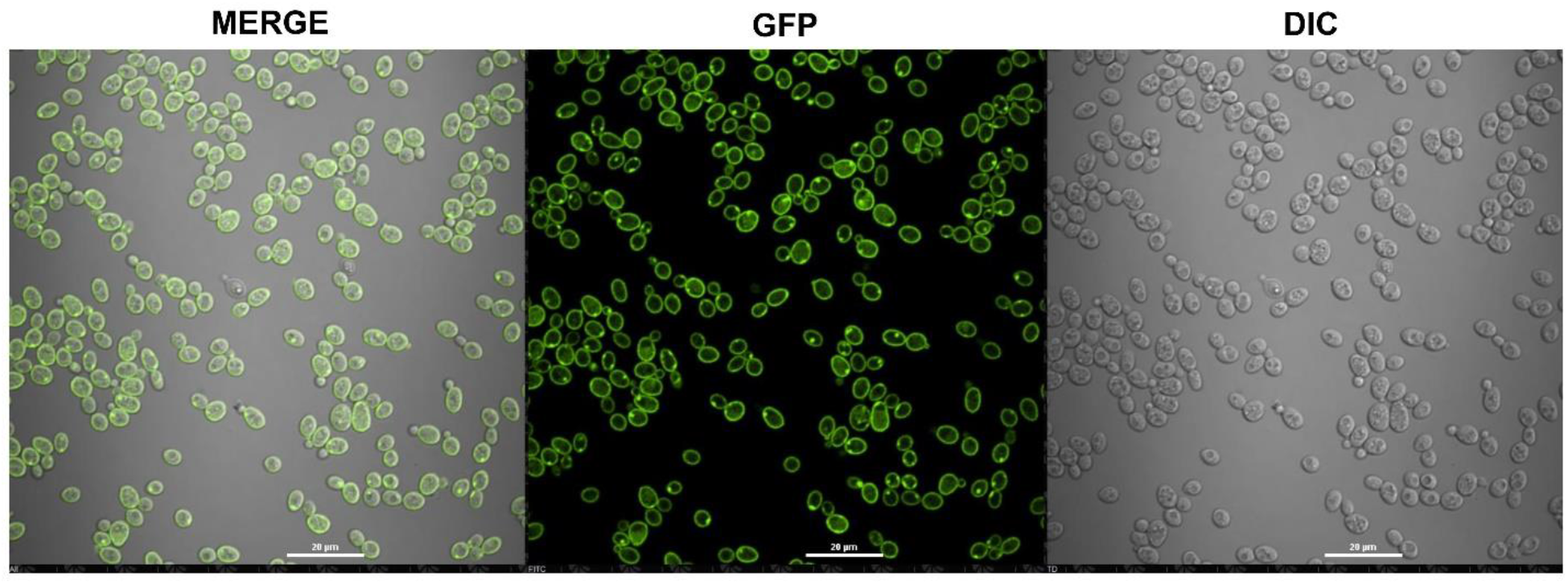
Confocal imaging of AD-*Cau*Mdr1-GFP strain. *S. cerevisiae* AD1-8u^-^ cells expressing GFP-tagged *Cau*Mdr1 protein at the plasma membrane.

### Nile red transport of *MDR1* knockout and overexpression strains

Given the alterations observed in the resistance attributes of the AD-*Cau*Mdr1-His strain, we next assessed the Nile Red efflux capabilities of the knockout and overexpression strains using a Flow cytometry-based accumulation assay. Nile Red is a known substrate of *Ca*Mdr1 and has helped contribute immensely to understanding the molecular mechanism of efflux by the transporter (Redhu et al., 2018). With this analysis, we not only planned to study whether Nile Red is also a substrate of *Cau*Mdr1, but also whether the observed increased resistance in the overexpression strain is indeed a function of its efflux. We evaluated the transport in the CBS10913T and *MDR1* knockout strain as well (**Figure 4**). As shown in **Figure 4A**, both CBS10913T and the *MDR1*Δ strain show similar levels of accumulated Nile Red which is evident from the statistically insignificant median fluorescent intensity (MFI) in the two strains. Contrarily, the host, AD1-8u^-^ strain and AD-*Cau*Mdr1-His strains displayed marked differences in the Nile Red retention levels, wherein the overexpression strain showed several-fold reduced Nile red fluorescence compared to the host (**Figure 4B**). Overall, it could be said that the transport assays corroborate our findings from the drug resistance studies.

**Figure 4:**
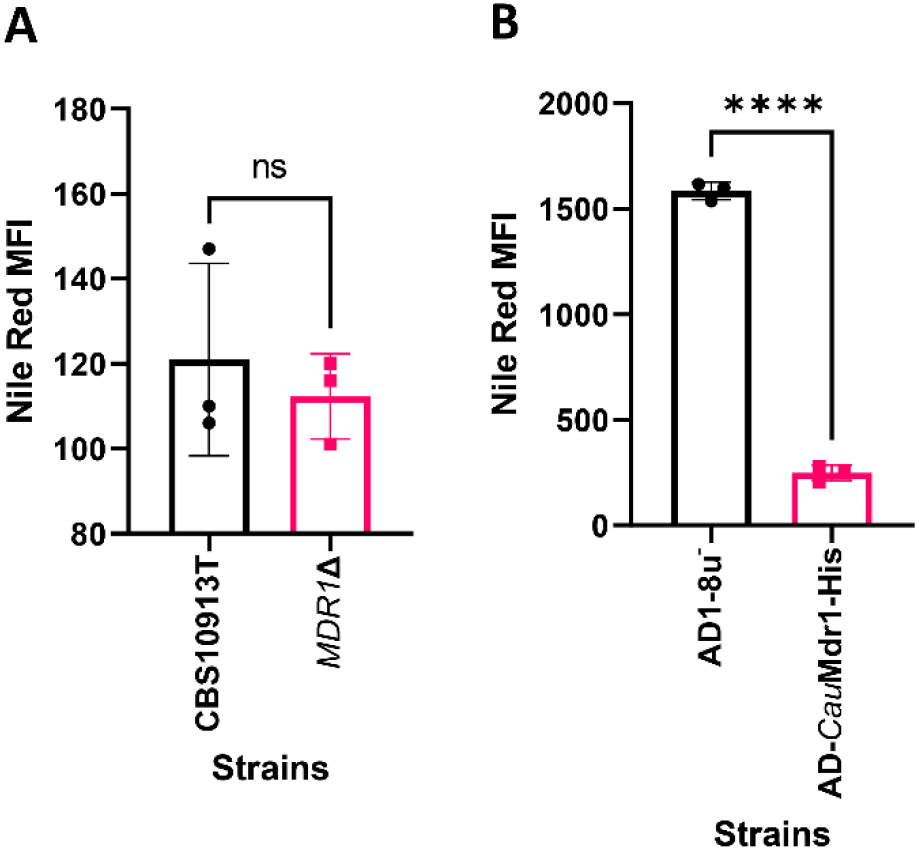
Whole cell-based substrate transport assay using Flow cytometry. Nile red accumulation in the **(A)** CBS10913T and the *MDR1Δ* strain **(B)** AD1-8u^-^ and the AD-*Cau*Mdr1 strain as represented by the median fluorescent intensity (MFI). The data are represented as mean ± SD. Statistical analyses were performed using the Unpaired T-test with Welch’s correction. ns represents non-significance whereas **** represents p-value <0.0001.

## CONCLUSIONS

The present study reports the first genome-wide inventory of DHA1 transporters in *C. auris*. We identified a total of 14 putative DHA1 transporters encoded in the genome. Interestingly, besides the ortholog of the prominent multidrug transporter *Ca*Mdr1, several orthologs of other transporters involved in the multidrug/multixenobiotic resistance phenomenon of yeast (such as the Qdr proteins) are present in its genome. Analysis of the transcriptomic dataset from the FungiDB also showed certain genes to be overexpressed in the biofilm stage, further strengthening their probable roles in the drug resistance phenomenon.

In the study, we also attempted to investigate the involvement of *Cau*Mdr1 in conferring resistance towards a panel of antifungal drugs/xenobiotics, some of which are previously shown to be substrates of Mdr1 and DHA1 transporters of other fungal/yeast species. While the deletion mutant of *MDR1* did not significantly alter resistance attributes of *C. auris* as reported before, overexpressing *Cau*Mdr1 in the heterologous AD1-8u^-^ system helped identify its several substrates namely, fluconazole, voriconazole, clotrimazole, ketoconazole, cycloheximide, 4-NQO, anisomycin, cerulenin, and Nile red. This study thus provides a platform for future structure-function studies on *Cau*Mdr1 and makes way for functional studies of DHA1 transporters in *C. auris*.

## FUNDING

This study is supported by funding from the Department of Biotechnology, Government of India to AB, RP, and AML through grant no. BT/PR32349/MED/29/1456/2019.

## ACKNOWLEDGEMENTS

The authors acknowledge the Central Instrument Research Facility at Amity University Haryana for confocal and Flow cytometry facilities. AB would like to acknowledge funding support from SERB, Government of India through the grant SRG/2019/000514. RK acknowledges JRF fellowship from SERB through the project SRG/2019/000514. SS acknowledges fellowship support from the ICMR, Government of India in the form of an SRF award.

## CONFLICTS OF INTEREST

None to declare

## Notes

### Competing Interest Statement

The authors have declared no competing interest.

