## Supplementary table for "Genomic landscape of the DHA1 family in the newly emerged pathogen *Candida auris* and mapping substrate repertoire of the prominent member *Cau*Mdr1"

**Supplementary table 1: List of strains used in the study**

| **S. No.** | **Strain ID** | **Description** | **Reference** |
| --- | --- | --- | --- |
| 1. | CBS10913T | Clinical isolate | (Wasi et al., 2019) |
| 2. | *MDR1*Δ | *MDR1* gene deleted from CBS10913T and replaced with NAT | This study |
| 3. | AD1-8u^-^ | MATα PDR1–3 ura3 his1 Δyor1::hisG Δsnq2::hisG Δpdr10::hisG Δpdr11::hisG Δycf1::hisG Δpdr3::hisG Δpdr5::hisG Δpdr15::hisG | (Lamping et al., 2007) |
| 4. | AD-*Cau*Mdr1-GFP | AD1-8u^-^, Δpdr5::pABC3-*Cau*Mdr1-GFP | This study |
| 5. | AD-*Cau*Mdr1-His | AD1-8u^-^, Δpdr5::pABC3-*Cau*Mdr1-His | This study |

**Supplementary table 2: List of oligonucleotides used in the study**

| **S. No.** | **Primer ID** | **Sequence (5'-3')** |
| --- | --- | --- |
| **Gene deletion and confirmation primers** | | |
| 1. | MDR1_P1 | GAGCCACAACTTCCCTATTCATTG |
| 2. | MDR1_P4 | CAGCCAAGGAAGCTACTGCCGCTAAC |
| 3. | MDR1_P5 | GGAGCACACTTCCGAAGATACGGCC |
| 4. | MDR1_P6 | GCGTCGACCTGCAGCGTACGATGTGGAGATTGAAGATGCGTTG |
| 5. | MDR1_P7 | CGACGGTGTCGGTCTCGTAGGATCCAAGTACGCTGGAGCAGGTG |
| 6. | MDR1_P8 | GTACAGTGAGCGGAGCCAACAATTTC |
| 7. | MDR1_P13 | GGTGTTTTCGCCGCTTTCGGAAAATGCC |
| 8. | MDR1_P14 | CAACAATAGCCATGGGAATGAAGAC |
| 9. | NAT_P2 | TGCGCACGTCAAGACTGTCAAGG |
| 10. | NAT_P3 | TGTGAATGCTGGTCGCTATACTGC |
| 11. | NAT_P9 | CGTACGCTGCAGGTCGACgccttccgctgctaggcgcgccgtg |
| 12. | NAT_P10 | GTCTACTACTTTGGATGATAC |
| 13. | NAT_P11 | TCTGTTCCAACCAGAATAAG |
| 14. | NAT_P12 | ctacgagaccgacaccgtcgggccgctgacGAAGT |
| 15. | MDR1_SEQ1 | GTCTGCACCCAAACAAAGACAATC |
| 16. | MDR1_SEQ2 | GGACATGCAACTGTCAAACGTAGAAC |
| **Cloning primers** | | |
| 1. | MDR1_PAC1 | CGCGATTAATTAAATGTTCCTCTATAAATTCGTCAGAG |
| 2. | MDR1_NOT1 | CGCGAGCGGCCGCAGGCACCTGCTCCAGCGTACTTGGAT |
